# A 3D Modeling Framework for Quantifying Variation in Soybean Root Structure Architecture

**DOI:** 10.1101/2025.05.06.652473

**Authors:** Joshua Carpenter, Sujata Bogati, Diane Wang, Jinha Jung

## Abstract

Root system architecture (RSA) underpins plant access to water and nutrients, making its characterization critical for improving crop performance in environments with limited soil fertility. However, current methods for quantifying root features face several challenges. They may rely on 2D images that suffer from occlusion, use expensive sensing technologies like X-ray computed tomography, or depend on 3D modeling approaches with assumptions about branching that make them difficult to generalize. To address these challenges, we introduce an open-source Python framework for quantifying RSA samples from 3D point clouds generated from low-cost photogrammetry. Critically, this method incorporates no assumptions about taxon-specific branching orientation, making it both well-suited for modeling naturally grown annual dicots such as soybean and generalizable across species. Using field-grown soybean as a test case, we demonstrate the utility of this framework to extract biologically meaningful 3D features of divergent root systems sampled across developmental stages and soil environments, and enable new analyses not possible with 2D approaches, such as modeling metabolic scaling relationships. Results indicate that, in our soybean samples, while certain individual features like taproot tortuosity are potentially influenced by the soil environment, and while roots in sandy loam exhibited greater feature plasticity, fundamental scaling properties remain consistent. By combining low-cost photogrammetry with 3D reconstruction of root systems from point clouds, this approach provides the plant science community with new opportunities for more comprehensive root studies.

## 1 Introduction

Plant roots are critical for the uptake of water and nutrients from the soil and for stabilizing the plant in its environment [Lynch, 1995]. The efficiency of these complex systems is determined by the spatial arrangement and physical characteristics of the root system [Herrero-Huerta et al., 2021], termed the root system architecture (RSA). Since RSA directly affects crop performance, there has been a growing interest in identifying and cataloging RSA traits to understand the arrangements that provide site-specific advantages for stabilizing crop performance. This is especially critical for improving food production in developing nations, where many agricultural regions are threatened by low soil fertility and water deficit conditions [Lynch, 2007]. However, comprehensively characterizing root system architecture for globally important crops, such as soybean (*Glycine max*), has been a major challenge.

Most conventional methods of phenotyping field-sampled root systems of crop plants rely on 2D imaging. These approaches are low-cost and relatively high-throughput. However, because of the complex 3D nature of RSA, features such as distribution, orientation, number of individual roots, and self-similarity are inaccurately captured and poorly analyzed with 2D methods, which suffer from occlusion [Wu et al., 2023]. RSA is a complex 3D object, so roots in the foreground appear to intersect with roots in the background when the object is projected onto a 2D imaging plane. Performing analysis of RSA in the native 3D allows these ambiguities to be resolved. 3D modeling further allows a whole range of new features to be extracted. In 2D, only major features can be sampled, and much of the architecture must be aggregated into physically ambiguous features, such as the number of crossing roots. 3D analysis enables access to the entire structure and the topology from the taproot down to the smallest root detectable in the 3D scan.

3D modeling technologies have recently been leveraged to great effect. Foremost among these methods are x-ray computer tomography (CT) imaging [Herrero-Huerta et al., 2021] and photogrammetric reconstruction [Dowd et al., 2021, Wu et al., 2023]. CT imaging has the advantage of being non-destructive – root systems can be sensed without removing the growth media. However, the technology is expensive, and plants must be grown in laboratory conditions [Dowd et al., 2021]. Photogrammetric methods, in comparison, are low-cost and achieve accurate 3D reconstructions of the RSA [Dowd et al., 2021]. Non-destructive photogrammetric imaging is possible if samples are grown in transparent media under laboratory conditions [Downie et al., 2012, Ma et al., 2019, Topp et al., 2013], while field-grown samples may be modeled if samples are exhumed [Okamoto et al., 2022, Wu et al., 2023]. While there is no one-size-fits-all method for modeling and analyzing RSA [Dowd et al., 2021], photogrammetric scanning offers the best low-cost, high-fidelity solution for measuring the RSA of exhumed samples.

Recently, a variety of methods have been developed to convert 3D point clouds of branching structures into models for parameter extraction. Point clouds are unstructured datasets, meaning the relation between points is not hard-coded into the data type. To the human eye, features in point clouds appear easy to discern, but to extract useful metrics from the RSA, methods must be implemented to recognize the constituent parts of the root from the point cloud and arrange them into the correct branching hierarchy. Most, if not all, approaches to 3D modeling of RSA are developed with specific modalities of raw data capture in mind. For instance, many methods have been developed for 3D modeling of RSA from CT volumetric images, such as RootForce [Gerth et al., 2021], Rootine [Gao et al., 2019], or RSAtrace3D [Teramoto et al., 2021]. There is some conceptual overlap between reconstruction methods designed for CT imagery and 3D point clouds. For instance, the idea of using level-sets to generate RSA skeletons using consecutive slices has been implemented across modalities [Mairhofer et al., 2016]. However, most methods are restricted to their mode of data capture. Some methods do exist which blend approaches, such as [Herrero-Huerta et al., 2021] in which tomato and maize root skeletons are extracted through laplacian contraction from CT images, and then branch diameters are extracted using TreeQSM [Raumonen et al., 2013], a method initially designed for forestry applications.

There are several approaches to modeling dendritic structures in 3D, originating in both agriculture and forestry applications. They can be grouped into the following categories: building-block, persistent-feature, and skeleton-first. A widely used building-block method is TreeQSM [Raumonen et al., 2013], such methods break a space into primitives and iteratively build structures from these primitives based on rulesets often influenced by a priori knowledge of the parameters constraining the dendritic structure [Hackenberg et al., 2014, Fan et al., 2020].

Persistent-feature methods detect structural objects in level sets of the data and then track these objects through successive layers to build topological skeletons. Examples include DIRT/3D which was designed for modeling maize roots from structure from motion (SFM) derived point clouds [Liu et al., 2021], and the work of Wu et al. in which rapeseed roots were grown in supported containers and modeled from SFM point clouds [Wu et al., 2023]. This method is ideal for monocot roots because of the general downward orientation of the systems, but fails for dicots with their orthogonal branching pattern and unpredictable directions.

Skeleton-first methods derive medial skeletons from the point cloud directly. These skeletons may be derived by a range of techniques such as using L1-median measurements to determine local centers of the point cloud [Huang et al., 2013], Laplacian contraction [Cao et al., 2010], or rotational symmetry [Tagliasacchi et al., 2009]. These skeletonization methods are popular in forestry applications using low-noise point clouds [Kim and Kantor, 2023] because they work well for structures like trees, which do not have a large disparity between the diameters of trunks and branches. In contrast, rooting systems where very fine roots branch directly off major taproots, as is the case with soybean root systems, such medial-seeking methods often omit small branches. In such scenarios, skeletonization based on graph analysis, such as the one presented in this paper, can perform well [Yang et al., 2024, Xu et al., 2007]. However, most existing models have been developed for tree modeling and incorporate assumptions about the branching patterns of trees [Yang et al., 2024] that may not be valid for root structures.

Despite the diversity of available methods described above, few are appropriate for the modeling of annual dicot RSA from 3D point clouds. Existing methods target monocot-specific architectures, are designed for 2D image or CT image input, or contain assumptions about branching structure that are often taxon-specific. To address the need for a 3D RSA modeling framework adapted for dicots, we present a method that generates highfidelity geometric twins of field-grown RSA samples collected from annual dicot species across developmental time. Using cultivated soybean as our test case, we (1) introduce an open source Python framework for quantifying field-grown RSA samples from point clouds, (2) design and identify 3D features that are sensitive to soil type, and (3) characterize variation in RSA across sandy loam and clay loam soils. This method of 3D reconstruction incorporates no assumptions about branching orientation and focuses on proper topology, enabling its generalizability across species and laying the groundwork for future development of 3D RSA feature databases of field-grown samples.

## 2 Materials and Methods

### 2.1 Data Collection

Soybean samples used in this study were grown during the Summer of 2022 at the Pinney Purdue Agricultural Center (PPAC) in Wanatah, Indiana, USA. Two sites within PPAC that offered different soil environments were used, hereafter referred to as the sandy loam and clay loam environments. Soil analysis results may be found in our companion paper [Bogati et al., 2025]. Plants were exhumed from both sites at three timepoints, corresponding approximately to growth stages V2-V6 (second to sixth trifoliate), V7-R2 (seventh trifoliate - corresponding to early flowering in most genotypes to full bloom), and R3-R7 (beginning pod to beginning maturity). These timepoints are designated as TP1, TP2, TP3, respectively. For details on root sampling methods, refer to [Bogati et al., 2025].

Close-range photogrammetric scanning was used to generate a 3D point cloud of 38 soybean root samples. For imaging, each sample was clamped into a holder and posed on a turntable, upside down in relation to its growing direction (see figure 1). The turntable rotated the sample while a Sony a7 MarkII DLSR camera captured photos every 5 degrees at a resolution of 5304px by 7952px. A small aperture (f/22) was maintained to create the largest depth of field possible during the image capture process. This allowed the lateral branches of each root to remain in focus even as they rotated towards and away from the camera position. The resolution and focus determine the size of the smallest structures that can be reconstructed. Roots with a diameter near the projected size of a single pixel will often be omitted. In our case, the projected size of a single pixel was approximately 0.05mm. After capture, Agisoft’s Metashape executed SFM 3D reconstruction. This method of reconstruction produced a scaleless point cloud of each root sample. Targets on the turntable were then used to scale the point cloud to the correct dimensions. The number of points varied widely by the size of the root. Approximately 200 thousand points represented samples from the earliest timepoint (TP1), while nearly 6 million points represented samples from the late season (TP3).

**Figure 1:**
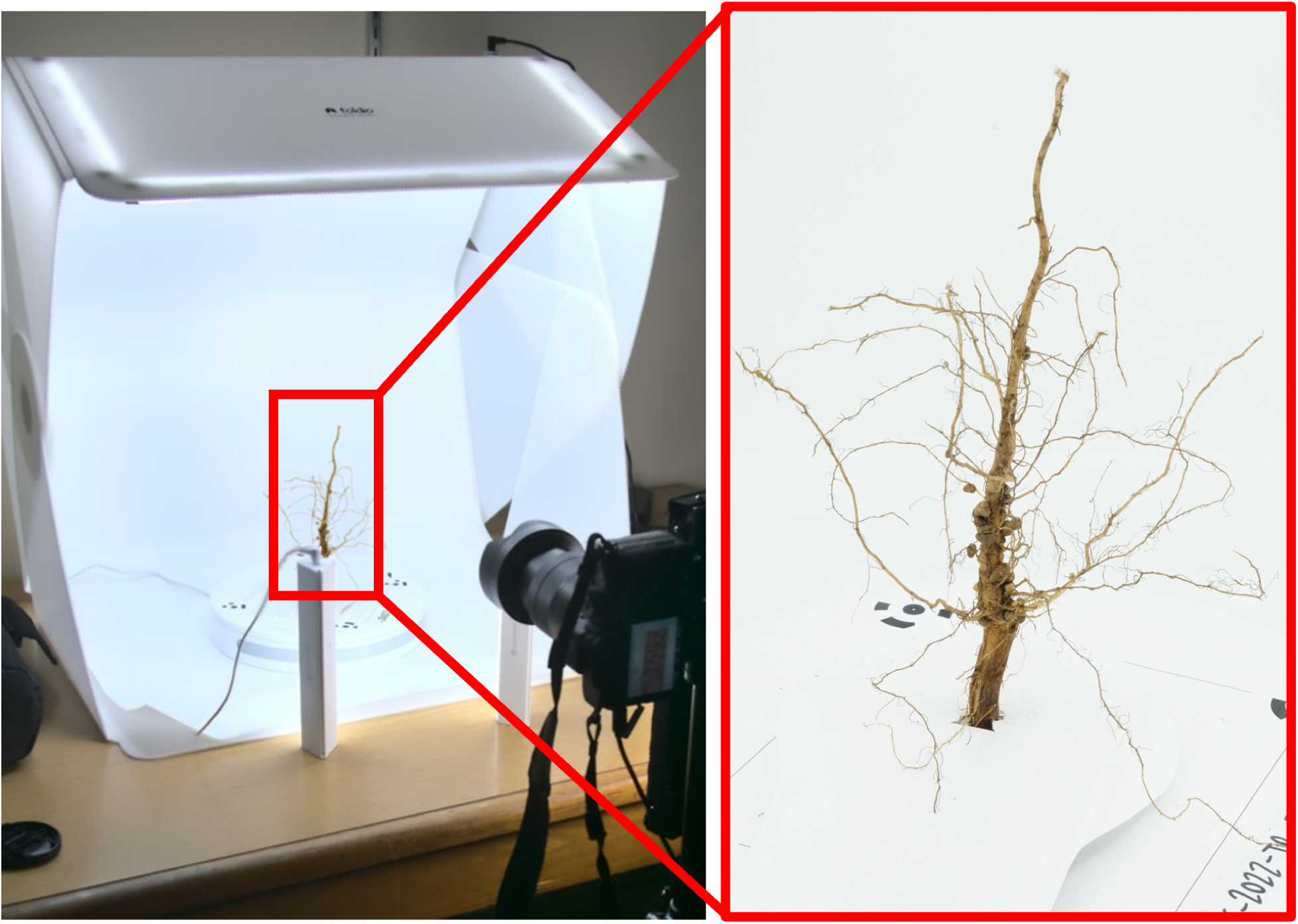
Close-range photogrammetric scanning booth with root. A root is clamped upside down in the booth and set on a turntable. The camera is stationary on a tripod while the sample turns.

### 2.2 3D Reconstruction of RSA

The full workflow for reconstructing root system architecture from point clouds is presented in figure 2 and is described in full below.

**Figure 2:**
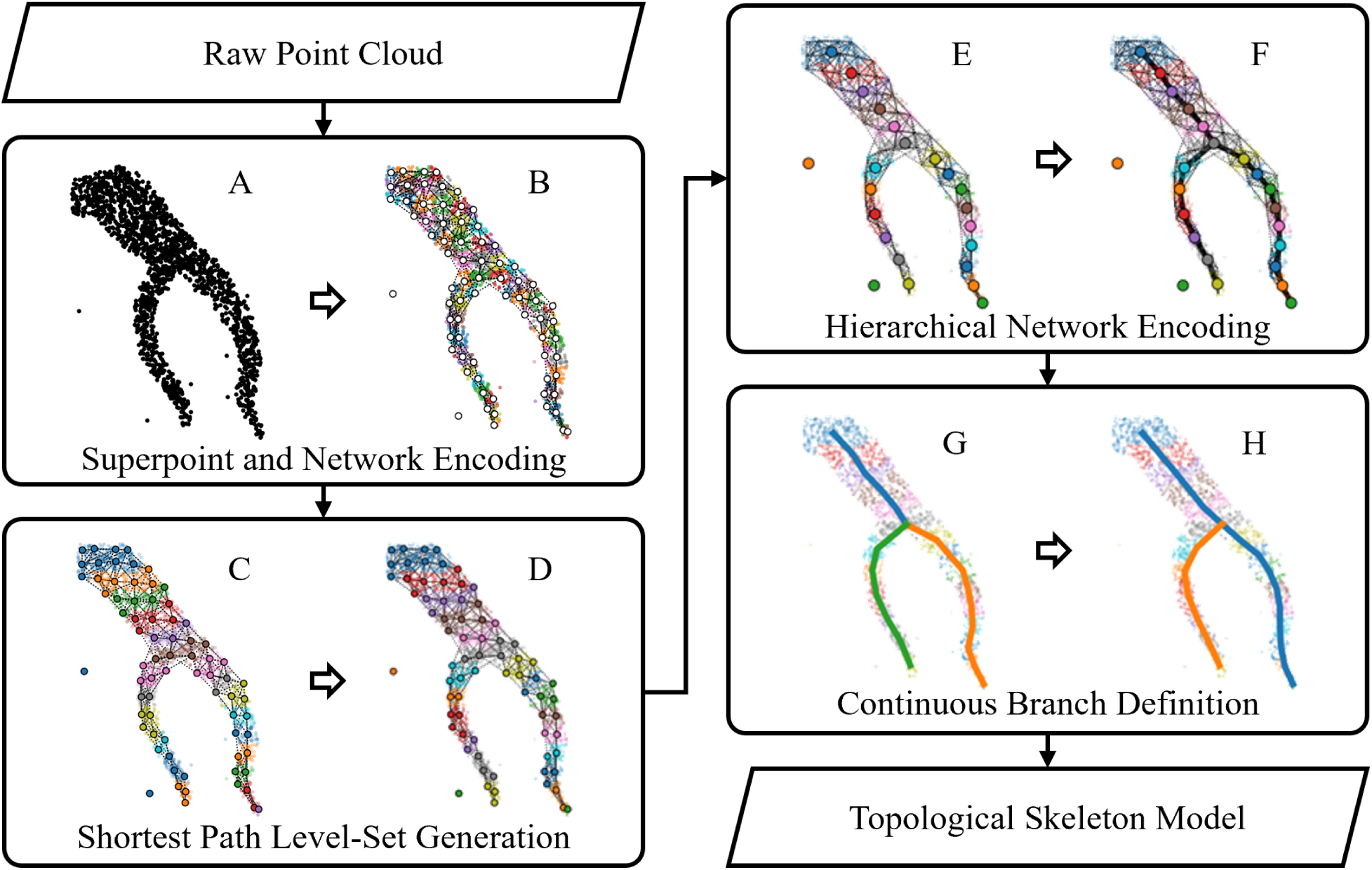
The modeling process flowchart. (A) Input is a raw point cloud. (B)*Superpoint and network encoding* creates small superpoints across the surface of the point cloud and defines connectivity. (C) *Shortest path level-set generation* bins superpoints based on their distance from the root base and (D) identifies unique clusters within the level-sets. (E)*Hierarchical network encoding* assigns a center to each block (F) to form a hierarchical network. (G) *Continuous branch definition* divides the network into ‘segments’ that link points of furcation then (H) uses Strahler ranking to determine root continuation.

*Superpoint and Network Encoding* starts by dividing the point cloud *P* into small neighborhoods - termed superpoints. Superpoints are geographically similar clusters of points that summarize the properties of a portion of a point cloud. Implementing superpoint encoding reduces the number of points within the input data, increasing processing speed and reducing sensitivity to point density variations in the input data. We divided the space into spherical superpoints of radius *r*. The radius should be chosen based on knowledge of the data. Gaps between points in *P* less than the radius will be ignored - so *r* should be conceptualized as the minimum point gap of importance. For this study, *r* = 0.5mm. This was achieved by first distributing temporary superpoint centers throughout the 3D space on an even voxel grid. The grid size is defined as 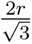 to ensure that a temporary center will fall within *r* of every point. The temporary center is then moved to the center of mass of all points falling within *r* of the center. This centering procedure evenly distributes centers on the surface of objects within the point cloud. These shifted centers become the centroid of the initial superpoint space *S*, and all points are then grouped into all superpoints that are less than *r* distant. This means that a single point may be a member of multiple superpoints. We developed a Python package, ShapeVoxelizer, to generate this specialized superpoint space. ShapeVoxelizer provides utilities for dividing point clouds into regular superpoint grids with cubic or spherical shapes, for shifting superpoints out of the standard grid arrangement for better representation of the point cloud’s geometry, for integrating multiple layers of nested superpoints, and for calculating descriptors of the points within each superpoint. [GITHUB LINK]

After building the superpoint space, the next step is to establish an initial network through the superpoint space. A network *N* consisting of nodes *S*, edges *E*, and weights *W*, which determine the cost of travel between superpoints. Since a set of initial superpoints *S* is only defined by their position in 3D space *S* ⊆ ℝ*^xyz^*, there is no inherently defined adjacency among the superpoints. We implement an adjacency definition that links superpoints that have a point in common. In other words, if there exists a point *p* such that *p* ∈ *s_i_* and *p* ∈ *s_j_*, edge *e_ij_* linking superpoint *s_i_* to *s_j_* is created. This method was first introduced by Raumonen et al. and was chosen because it makes the network more sensitive to gaps in the surface of the original point cloud [Raumonen et al., 2013]. The costs of travel along the edges *E* are defined by the cost function *W* : *E* → ℝ, which maps any edge *e_ij_*∈ *E* to a single cost value calculated as

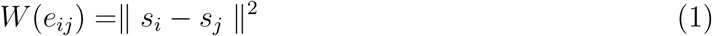

the square of the Euclidean distance between two adjacent superpoints *s_i_*∈ *S*. Squaring the Euclidean distance encourages the least-cost routing routine to choose routes with smaller gaps between nodes and discourages the selection of longer, more direct routes [Carpenter et al., 2022]. For building this network, and for building and extracting features from hierarchical networks, we developed the package SuperpointGrapher as a wrapper for the Python package Networkx [Hagberg et al., 2008]. Figure 2.B illustrates the results of the superpoint and network encoding step.

*Shortest Path Level-set Generation* is the next step in our RSA reconstruction. The goal of level set generation is to divide the set of superpoints *S* into clusters that each describe a narrow cross-section of a single root structure. Three procedures are implemented to achieve this: shortest path calculation, distance binning, and connected component analysis. This portion of the procedure closely follows that presented in [Xu et al., 2007].

The shortest path from each superpoint *s* ∈ *S* to the base of the root structure was calculated through the network *N*. We used Dijkstra’s shortest pathing algorithm. Using these distances, each superpoint is placed into a bin based on the distance from the superpoint to the base of the root structure. As was done in [Yang et al., 2024], we reduce the widths of the bins toward the branch tips of the branches. The initial bin width is set at 10*r* and reduces linearly to 2*r* at the branch tips. This ensures that the bins are scaled according to the size of the initial set of superpoints. We designed our modeling method to be agnostic to scaling properties of the dendritic structure being modeled. For this reason, we do not use an exponential scaling function to reduce bin widths. Figure 2.C shows level-set creation. Distance binning recognizes the distance from the base of the root structure, but does not divide individual roots into separate clusters. This can be seen in figure 2.C where points comprising different branches of the root are grouped into the same level-set. To discriminate between separate branches, the network *N* was subset for each level-set of superpoints. Then, connected component analysis was performed on the subset network using the functionality of the Networkx package. This divided each level set into clusters based on network connectivity. The result of this processing is shown in figure 2.D

*Hierarchical Network Encoding* begins with the creation of a new superpoint space *B* that converts each level-set cluster into a new superpoint, which we will refer to as a ‘block’ to differentiate from the small superpoints generated earlier (Figure 2.E). Because of the preceding steps, we can now conceptualize all the points constituting each of the new superpoints *B* as representing the surface of a cylinder. These blocks will form the base unit in the root reconstruction. The centroid of each block *b* ∈ *B* is the center of mass of the constituent superpoints. This center is assumed to fall near the central thread of the represented root structure.

A new network *H*, consisting of nodes *B* and edges *C*, encodes the connectivity between the blocks. The original superpoint network *N* is used to determine which blocks are connected. Blocks are comprised of many superpoints from the initial superpoint space *S*. So, if there is a superpoint (*s_a_* ∈ *b_i_*) within a block *b_i_*, having an edge (*e_ab_*) with another superpoint (*s_b_* ∈ *b_j_*) belonging to an adjacent block *b_j_*, then the two blocks *b_i_* and *b_j_* could potentially be connected with edge *c_ij_* in the new network *H*. However, not all connections are used to build network *H*. To ensure that the resulting network *H* represents the root structure, each block is connected to only one other block whose distance to the root base is less than itself. Figure 2.F shows the connections of the network *H*. This step effectively forms the skeleton of the root system.

*Continuous Branch Definition* defines how segments are arranged into branches. The network through the blocks forms a tree structure - it is non-looping. From this tree, individual branches will be defined. The first step is to build a hierarchical model from the skeleton, where blocks are grouped into segments (Figure 2.G). These segments are then arranged in parent-child relationships. Each segment represents a string of blocks linking one point of furcation with the next. Figure 3.A shows an example of this hierarchical structure.

**Figure 3:**
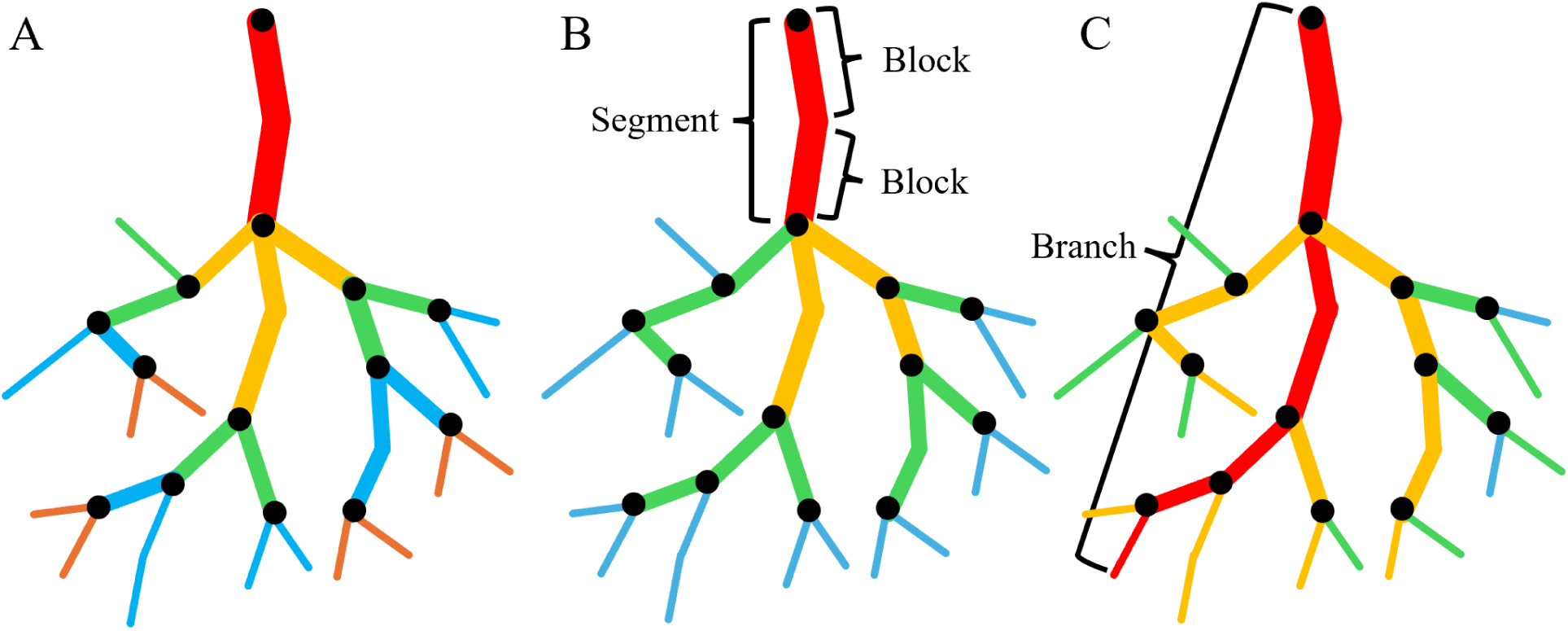
Various options for ranking a hierarchical tree structure. A) Top-down ranking. Each point of furcation changes the rank of all child branches. B) Strahler ranking. Rankings are based on topology, and similarly sized segments are ranked similarly. C) Contiguous branch recognition.

We then ranked each segment based on the Strahler method. Strahler ranking has long been used in studies of dendritic structures as a way to identify the hierarchical order of a particular branch while ensuring that branches of the same ranking have similar sizes [McMahon and Kronauer, 1976, Strahler, 1958]. The tips of each branch are given a rank of 1. Where two segments of the same rank join, the parent segment’s rank is increased by 1. Where two segments of differing ranks join, the parent segment takes the rank of the larger of the two child ranks (See figure 3.B).

Finally, continuous branches were defined using the Strahler ranking. Determining the continuation of a root segment through points of furcation requires a rule set for determining which child is a continuation of the parent segment. Figure 3.C gives an example of structure continuation. Continuation rules that used the diameter and pointing direction of segments were tried, but we found these rule sets to be unreliable since blocks near a point of furcation may include points from both child branches, causing the diameters and pointing directions to have more error close to the point of furcation. Therefore, we developed a rule set based on the more stable metrics of the topology: at a point of furcation, the child segment with the highest Strahler rank is determined as the continuation of the parent; while, in the case of two children of equal rank, the child with the longest cumulative length (the sum of the lengths of all network connections upstream of the point of furcation) would be identified as the continuation of the parent. Results are shown in figure 2.H. Figure 4 shows the full process, from point cloud to cylinder model, implemented on one of the soybean samples.

**Figure 4:**
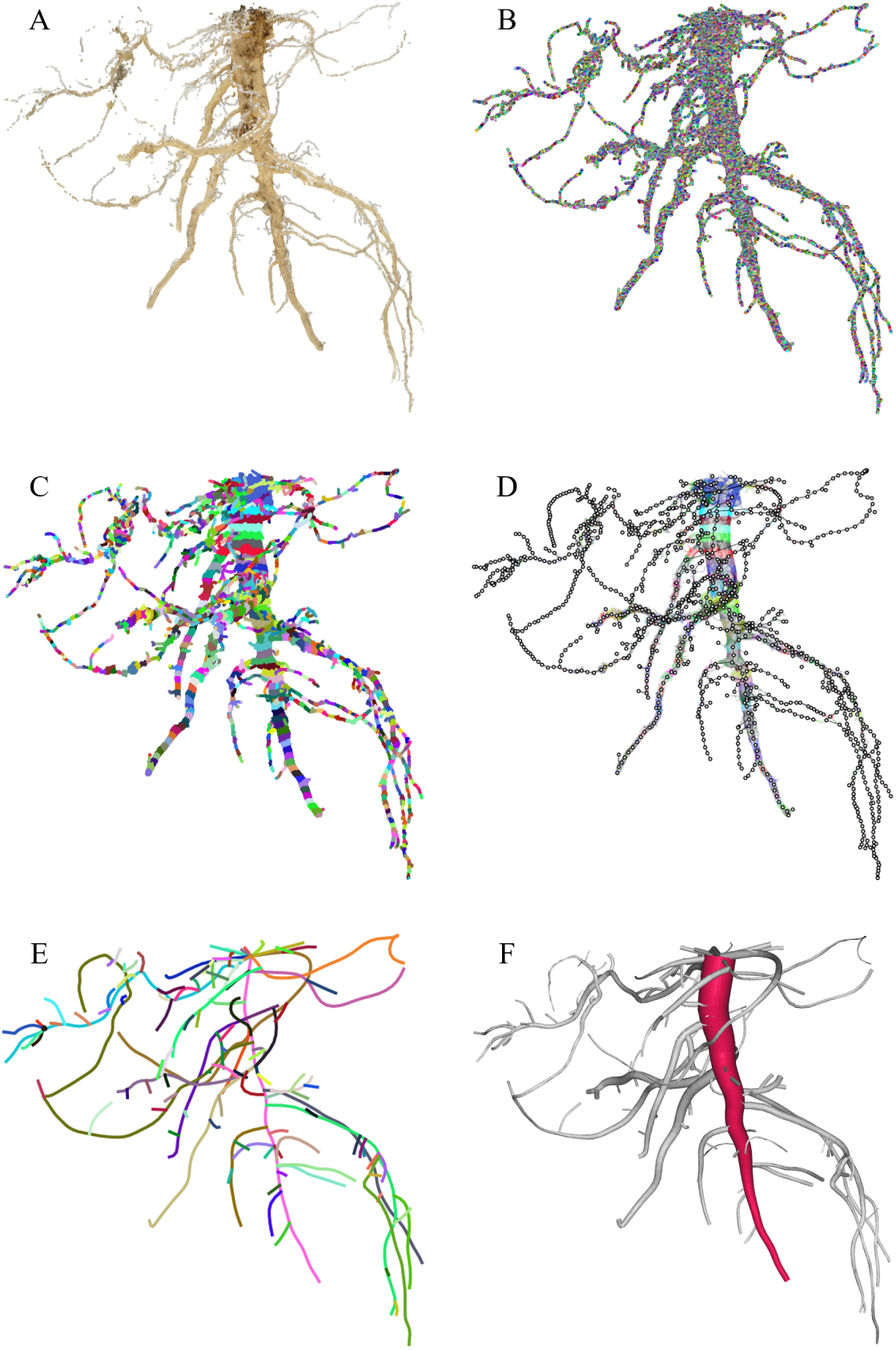
The modeling process demonstrated on a root sample. A) The raw point cloud. B) The result of initial *Superpoint and Network Encoding*, points colored by superpoint membership. C) The result of *Shortest Path Level-set Generation*, points colored by block membership. D) The result of *Hierarchical Network Encoding*, points colored by block membership. E) The results of *Continuous Branch Definition*, branch definitions indicated by color. F) The resulting 3D model of the RSA. The taproot is shown in red, demonstrating the encoded topology.

### 2.3 Outputs of the Topological Skeleton Model

The process laid out above creates a hierarchical skeleton of the RSA. Encoded in the skeleton is the relationship between each branch, segment, and block with the constituent points in the point cloud. This structure allows features to be calculated for individual blocks and then aggregated and analyzed at the segment or branch level, while considering branching topology.

Calculating branch lengths and diameters is necessary for subsequent phenotypic analysis. For a given block, we define the radius of the cylinder as the average of all the normal distances between the points comprising a block and the root skeleton. A moving window of 3 blocks was used to smooth the radii to reduce the influence of outliers. To visually check that the radii calculated for each block accurately model the point cloud, we generated 3D models of each root system modeled in this study based on the hierarchical skeleton and block-level radii. Figure 4.F shows an example with the tap root highlighted to demonstrate the hierarchical topology embedded in the reconstruction of the root system.

Another output of this workflow is a hierarchical classification for all points in the original root point cloud. Figure 5 shows the point cloud of a specific sample in which each point has been colored based on the branch it has been used to model. This segmentation allows other modeling methods to be easily integrated into the workflow presented in this paper.

**Figure 5:**
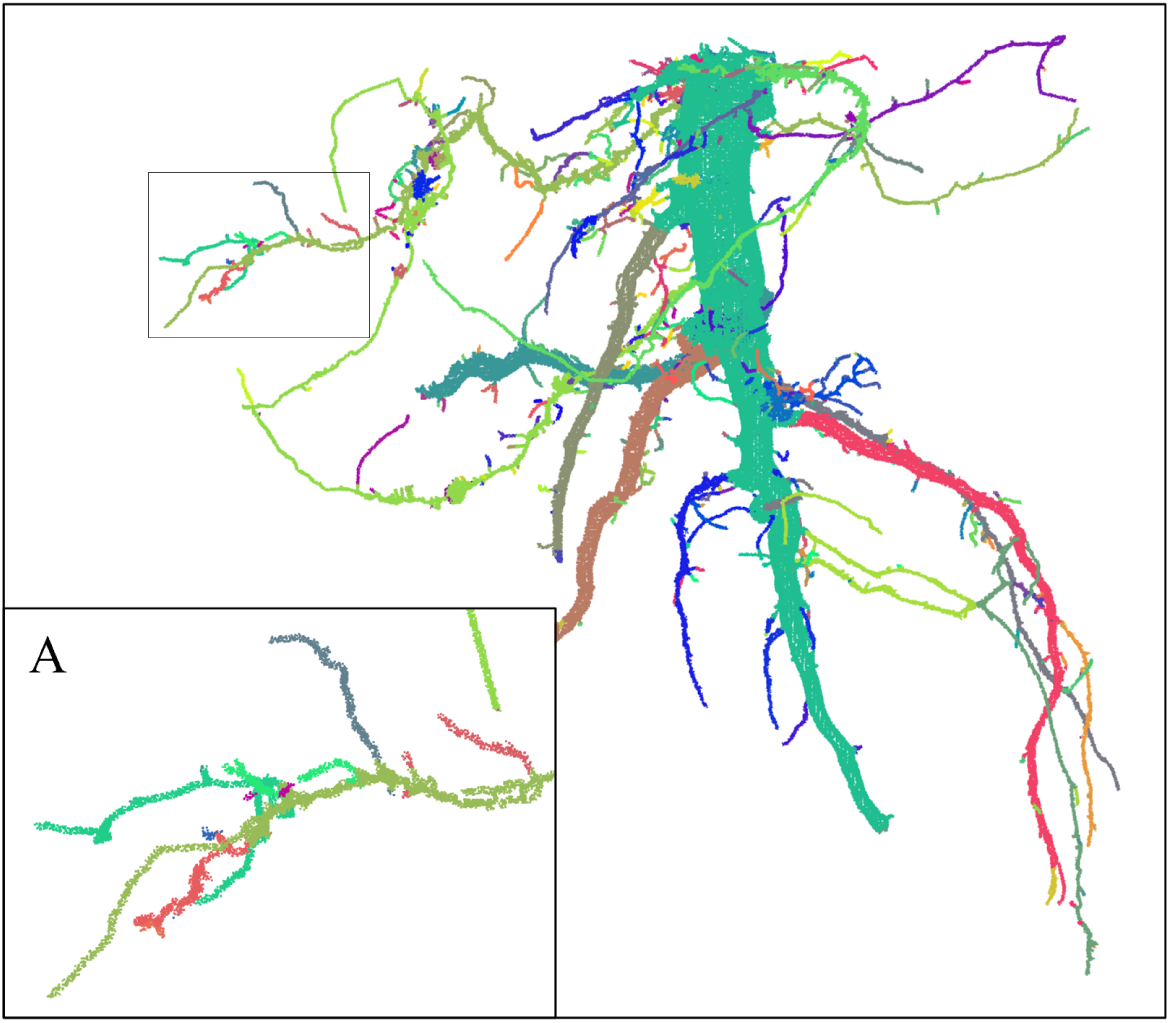
The point cloud of a root sample with each point labeled based on its branch membership. A) A close-up of a part of the point cloud.

### 2.4 RSA Analysis

#### Geometric Model Accuracy

After 3D modeling, the samples are analyzed to determine the accuracy of the 3D models. The geometric accuracies of the 3D RSA models were evaluated by observing the degree to which the skeleton and associated branch radii explain the positional variation in the point cloud. To do this, the residual for each point in the point cloud was calculated as the normal distance between a point and the surface of the cylinder describing its associated block. The significance of the size of a residual is dependent on the size of the feature being modeled, so the ratio between the residual and the diameter of the branch associated with the residual was additionally computed.

To identify extractable 3D features sensitive to soil type, and to characterize variation in RSA across sandy loam and clay loam soils, several analysis techniques were employed to study the features extracted from the reconstructed RSA.

#### Metabolic Scaling

The first metabolic scaling metric assessed was area preservation. Area preservation tendencies of roots grown in clay loam and roots grown in sandy loam were computed using the cylindrical model of the RSA. The area ratio *a_r_* is calculated between each parent-child pair as:

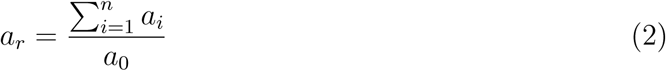

where *a*_0_ is the cross-sectional area of the parent branch, and the numerator is the summation of the cross-sectional areas of the *n* child branches.

Next, self-similarity was quantified by analyzing the scaling exponents of branch diameter and branch count across Strahler orders. Since our 3D RSA models encode the Strahler ranking of each branch within the network, we count the number of root segments and average diameter of the root segments within each Strahler rank. These values are plotted in figure A2 on a semi-log plot (with the *y*-axis in logarithmic scale and the *x*-axis as Strahler rank) for all samples in each timepoint. In these plots, the slope of the line represents the exponential rate at which these properties change with increasing branch order. We will refer to these metrics as the diameter ratio *R_d_* and the branching ratio *R_n_* calculated as a function of their rank *k* by the following equations:

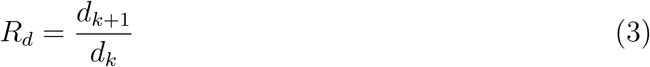

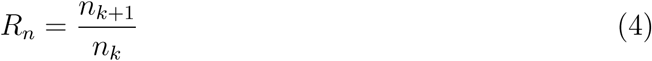

#### Feature Significance

The separability between the distributions of the sandy loam and clay loam samples was quantified using Fisher’s discriminant score, given by

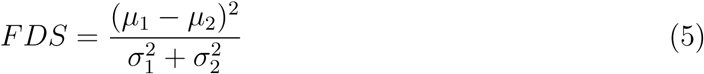

where *µ*_1_ and *µ*_2_ are the means of the two distributions, and 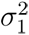 and 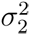 are their variances. The statistical significance between the feature means was assessed using the two-sample t-test. Finally, the statistical significance between the variation of the feature distributions was assessed using Levene’s f-test.

#### Feature Space Analysis

The significance of individual features may not be evident when analyzed in isolation, but can emerge through their interactions with other features. To better understand the structure of the feature space, we apply Principal Component Analysis (PCA) to identify combinations of features that capture the greatest variance in the data. Additionally, we use Linear Discriminant Analysis (LDA) to find feature combinations that maximize separability between responses associated with different soil types.

## 3 Results and Discussion

In total, 30 soybean root systems were sampled and reconstructed in 3D. These samples span three developmental timepoints and two soil environments. Examples of these models are shown in figure 7. From each RSA sample, 39 biologically-meaningful features were calculated (Table 2) and analyzed to identify features sensitive to soil variation and to describe the RSA response to the soil environment.

### 3.1 Geometric Model Accuracy

The root-mean-square error of the residuals between skeleton/branch radii and positional variation in the point cloud was 1.1mm, with an average residual of −0.1 ± 1.1mm. The full distribution of the residuals is shown in figure 6.A. This attests to the accuracy of our geometric RSA models and the precision of the source point cloud. Figure 6.B shows the error ratio between the residual and branch diameter for each point. The inflection point in the error ratio curve (about 2-3mm in figure 6.B) aligns with the lower limit of our block size. In our 3D models, the block size tapers from 10*r* (5mm) at the base of the root system down to 2*r* (1mm) at the tips. Since twists and turns in the root path smaller than the block size cannot be captured, the error ratio increases for the smallest roots.

**Figure 6:**
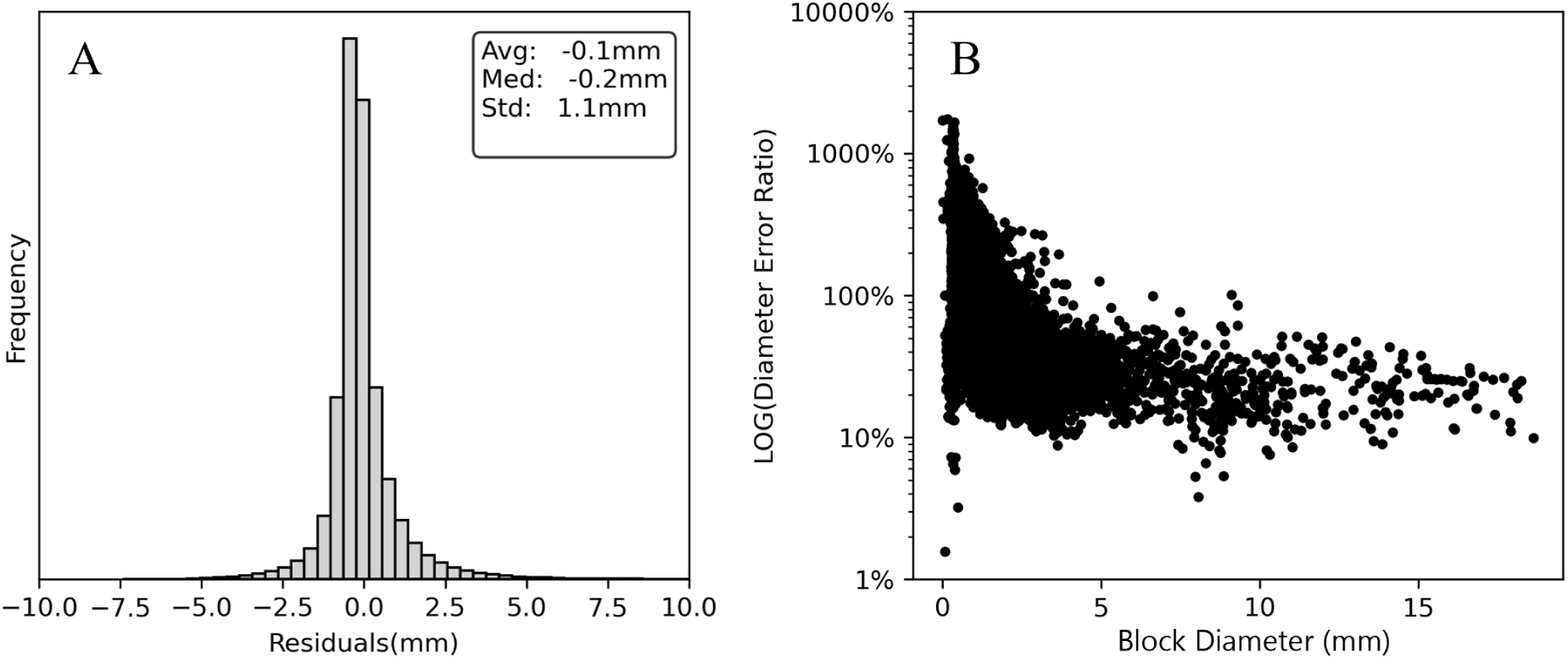
The distributions of select features.

**Figure 7:**
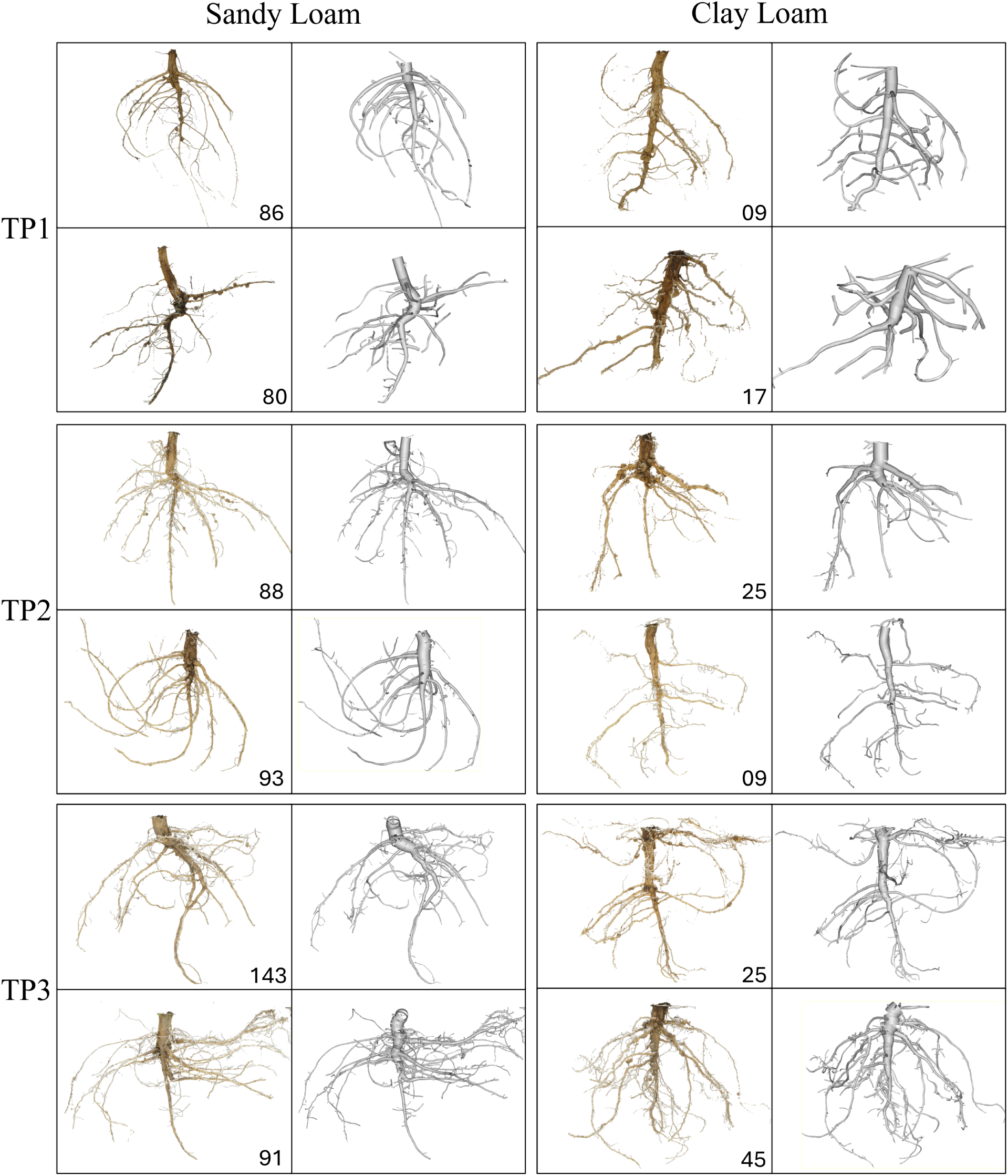
The point cloud and derived hierarchical cylinder model of 2 samples from each of the 6 experimental units.

### 3.2 Metabolic Scaling

3D RSA models enable the extraction of biologically-meaningful information such as metabolic scaling relationships, which refers to how biological processes such as nutrient transport, and structural integrity scale with size. Root architecture is directly affected by the metabolic cost of vascular transport efficiency and structural optimization, since these functions alter the number and length of roots that can be produced and sustained [Lynch, 2022]. We therefore analyzed metabolic scaling relationships between every parent and child pair within each sample, focusing on (1) area preservation and (2) self-similarity.

#### Area preservation

Cross-sectional area is a key structural indicator of variation in growth or nutrient acquisition strategies, since hydrodynamic resistance depends on the radius of the vascular structure [Bentley et al., 2013]. As far back as Leonardo da Vinci, it has been hypothesized that in an optimal vascular branching system cross-sectional area will be preserved throughout the branching structure; that is to say, the cross-sectional area of the parent branch will equal the sum of the cross-sectional areas of its children [McMahon and Kronauer, 1976].

In 3D, the degree to which a root system meets this area preservation benchmark can be directly and exhaustively measured. Area ratio was first developed for studying tree structure architecture. Recent work on tree structure has shown that the average value of this ratio is expected to be between 0.90 and 1.05, but varies by tree species and environment [Bentley et al., 2013, Eloy et al., 2017]. Our analysis indicated that the mean area ratio for our soybean samples (1.04 for Sandy Loam and 1.05 for Clay Loam) fell within the expected value range set by previous research into tree structures (figure A1). This suggests that similar metabolic and structural efficiencies control both tree structure architecture and the variation in RSA. This finding contrasts with recent work, which suggests that metabolic scaling rates differ significantly between root systems and other above-ground plant structures, such as trunks and branches, that have a supportive functionality [Sopp and Valbuena, 2023]. While future studies will be required to definitively show if root structures and above-ground dendritic structures follow differing metabolic scaling rates, our results demonstrate that features extracted from our 3D modeling procedure accurately reflect properties of natural branching structures. When comparing samples across soil environments, we found no significant difference in area preservation, indicating that this property is largely independent of soil type.

#### Self-similarity

Self-similarity reflects an underlying optimization principle, where branch diameter and segment count follow predictable scaling laws to balance mechanical stability (anchorage in the soil), hydraulic conductivity, and metabolic cost. Across all timepoints, the rate of diameter increase consistently fell within a window of 1.95–2.05 for both clay loam and sandy loam environments, meaning the diameter of most roots doubled at each change in strahler rank. A notable exception to this trend is seen in TP2 where clay loam samples appear to have a lower rate of diameter increase compared to their sandy loam counterparts (figure A2). However, given that TP1 and TP3 do not show this trend, we are inclined to suggest that this observation is an anomaly in the samples used to study TP2. Similarly, no significant difference was found in the branching rate within the two environments. Taken together, these results suggest that soil texture did not have a major effect on the self-similarity of the RSA in these soybean samples.

### 3.3 Feature Significance

We next examined which 3D RSA features had the greatest differences between samples grown in sandy loam versus clay loam environments. Results from the Fisher’s discriminant, t-test, and f-test are summarized in figure A4. From this analysis, we identified nine features of interest: the length of the taproot (*length*), the volume of all the root mass in the sample (*cumul volume*), the diameter of the fattest part of the taproot (*diameter MAX*), the average diameter of the tap root (*diameter AVG*), the straightness of the taproot (*tortuosity*), the number of branches (*branch count*), the number of furcation nodes (*node count*), and the rate of increase in diameter between strahler ranks (*diameter increase ratio*). The distributions of these nine features in both clay loam and sandy loam are shown in figure A3.

These analyses of our RSA models show that samples grown in sandy loam soil appeared more developed than their clay loam counterparts at the same timepoint. The sandy loam samples had longer tap roots in TP2 and TP3 indicative of slower growth rates (figure A3.A), had more structural volume (figure A3.B), reached deeper into the soil (figure A3.C), and had more branches than clay loam samples. While samples grown in clay loam tended to have tap roots with thicker diameters throughout their length (figure A3.E) with high tortuosity (figure A3.F) and fewer branches (figure A3.G).

#### Sandy Loam

Samples grown in sandy loam showed more feature plasticity in both size and architecture complexity. figure A4 shows that the cumulative volume (*cumul. volume*) in TP3 is statistically different between the two environments and highly separable, initially indicating that clay loam reduces root growth. Comparing the distributions reveals that one soil is not explicitly increasing or reducing RSA size. Rather, the variability in the cumulative volume of sandy loam roots indicates that the soil environment is controlling the potential range of length and diameter available to the rooting system. This variability in size is further seen in the depth reached by the rooting systems (*depth span*), where sandy loam roots show both better soil penetration and a high degree of variation in depth between samples.

Further highlighting the greater plasticity in RSA features possible to sandy loam samples are the features *branch count* and *node count*. Sandy loam samples tend to have more branches and more furcation nodes than their clay loam counterparts. However, these trends are not statistically significant, nor are the distributions of these features separable between the environments (figure A3). However, comparison of the distributions shows the impact of the sandy loam environment on the RSA. These features have a significantly higher variability in sandy loam, as indicated by the f-test of the variance of the *branch count* feature. The sandy loam samples have far more variability in both size and bushiness than those samples grown in clay loam. This variability suggests that exploratory potential is greater in sandy loam. Roots are free to grow longer and generate more laterals to secure ample nutrients and water.

#### Clay Loam

In contrast with the high variability of the sandy loam RSA features, samples grown in clay loam are generally smaller and more consistent in their structural composition. The average diameter of the tap root (*diameter AVG*) is an exception to the trend. Early in the growing season, the average diameter of clay loam taproots is significantly smaller than sandy loam samples. However, as growth progresses, our finding shows that the average diameter of clay loam taproots first matches and then exceeds the average diameter of sandy loam samples by the late-stage growing (figure A3.E). This pattern is not, however, matched by the maximum taproot diameter. Nor does taproot length correlate with average diameter. For instance, at TP1 when clay loam and sandy loam taproots are the most different in average diameter, their lengths (fig SA3.A) are nearly identical. This seems to indicate that clay and sandy loam samples have nearly the same depth penetration rate; however, the sandy loam allows the tap root to thicken earlier, while the clay loam slows the root’s thickening.

This indication of slower rate of tap root thickening is supported by the comparison of tortuosity. In this work, tortuosity is calculated as the straight-line distance between the base and tip of the tap root, divided by the distance by skeleton between the base and tip of the tap root. A tortuosity value of 1 indicates a perfectly straight root. Lower values indicate curvature in the taproot. The distributions of tortuosity are shown in figure A3.F. The plot indicates that clay loam samples are significantly more tortuous (twisted) than their sandy loam counterparts. This conforms with the finding that clay loam restricts the thickening rate of the tap root. The tap roots of both sandy and clay loam samples penetrate the soil at the same rate, but clay loam samples have thinner roots. Thinner roots conform more readily to local variation in soil density, resulting in a tap root which is more tortuous.

### 3.4 Feature Space Analysis

To capture the feature interactions that account for the greatest variance in the data, we performed PCA. The results of our PCA analysis are shown in figure A5. The first principal axis (PC1) explains 49% of the variance, and, unsurprisingly, features that capture the size of the root contribute the greatest. The second principal axis (PC2) explains 11% of variance and is driven by *percent of cumul. volume* (10%), *diameter AVG* (8%), *percent of cumul. length* (7%), *internodal length AVG* (7%), and *rate of diameter reduction* (7%). Visually, PC2 seems to represent the variation between soil types. The features with the largest loadings in PC2 are those features that together span the potential variability caused by soil type.

LDA gives a complementary view of the data. While PCA reveals those features that interact to find the axis of the greatest variance, LDA reveals those features whose interactions describe the axis of greatest separability. Results of LDA analysis are shown in figure A6. The first discriminant axis explains 79% of separability with volumetric features such as *volume*, *cumul. volume*, and *average 2D convex hull area* being the largest contributors.

The second discriminant axis explains 14% of the separation, with features describing the depth (*depth span*, *total depth span*, and *length*) and branching rates (*node count* and *branch count*) being most impactful.

### 3.5 Key Findings

#### Model Accuracy

We developed a high-resolution, generalizable 3D modeling framework tailored to the complex RSA of annual dicots. Unlike previous methods constrained by imaging modality or designed for monocot structures, our pipeline accurately reconstructs field-grown dicot RSA from point clouds without imposing assumptions about branching orientation or topology. The accuracy of our models is demonstrated in several ways. First, the geometric models are highly precise, with an average residual of −0.1 ± 1.1 mm and a root-mean-square error of 1.1 mm, enabling the extraction of fine-scale features, down to 1 mm in diameter, even in densely branched systems. Second, our models preserve the topological and geometric detail necessary to quantify biologically meaningful variation, as reflected in the metabolic scaling analysis showing consistency with natural dendritic patterns. Finally, these high-fidelity reconstructions detected phenotypic responses to soil type in soybean roots, confirming that our method captures natural RSA variability and supports downstream phenotyping aimed at improving crop resilience across environments.

#### Parameters of Importance

Several methods were employed to identify features that exhibit environmentally induced variation. Separability of clay and sandy loam RSA features was determined using discriminant analysis; statistical significance between feature means was measured using the t-test; differences in feature variance across soil types were analyzed with the F-test; and interactions among features describing variation and separability in the feature space were examined using PCA and LDA loadings. The key features identified by this paper and the test which caused them to stand out as a feature of importance are summarized in table 1. Different groups of features capture distinct aspects of RSA variation driven by soil conditions. Some of the most powerful variables for separation are those that relate to the length and shape of the taproot (*length*, *tortuosity*, *diameter increase ratio*, and *diameter MAX*). Features that describe the number of branches and branching rate (*branch count* and *node count*) have variances which are affected by soil type and can discriminate between impacted RSA when linearly combined with depth-related features such as *depth span*, *total depth span*, or *length*. Finally, though they do not show much separability between clay and sandy loam impacted RSA, features related to ratios between taproot volume or length to the total volume or length of the rooting system seem to serve as parameters that span the space of feature variability.

**Table 1:**
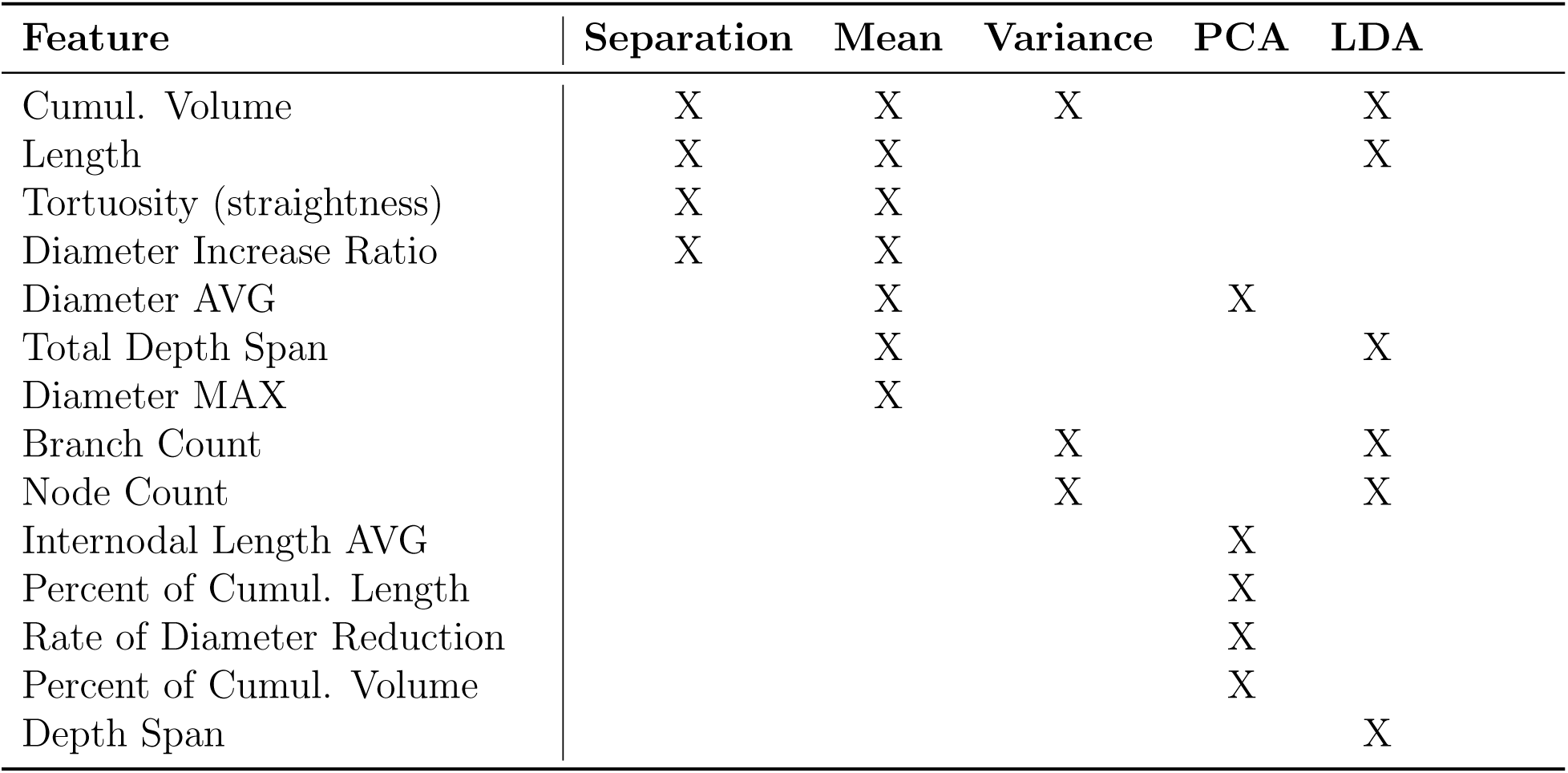
A summary of the features identified by this study that capture the potential variation in RSA due to clay or sandy loam soils.

| Feature | Separation | Mean | Variance | PCA | LDA |
| --- | --- | --- | --- | --- | --- |
| Cumul. Volume | X | X | X |  | X |
| Length | X | X |  |  | X |
| Tortuosity (straightness) | X | X |  |  |  |
| Diameter Increase Ratio | X | X |  |  |  |
| Diameter AVG |  | X |  | X |  |
| Total Depth Span |  | X |  |  | X |
| Diameter MAX |  | X |  |  |  |
| Branch Count |  |  | X |  | X |
| Node Count |  |  | X |  | X |
| Internodal Length AVG |  |  |  | X |  |
| Percent of Cumul. Length |  |  |  | X |  |
| Rate of Diameter Reduction |  |  |  | X |  |
| Percent of Cumul. Volume |  |  |  | X |  |
| Depth Span |  |  |  |  | X |

**Table 2:**
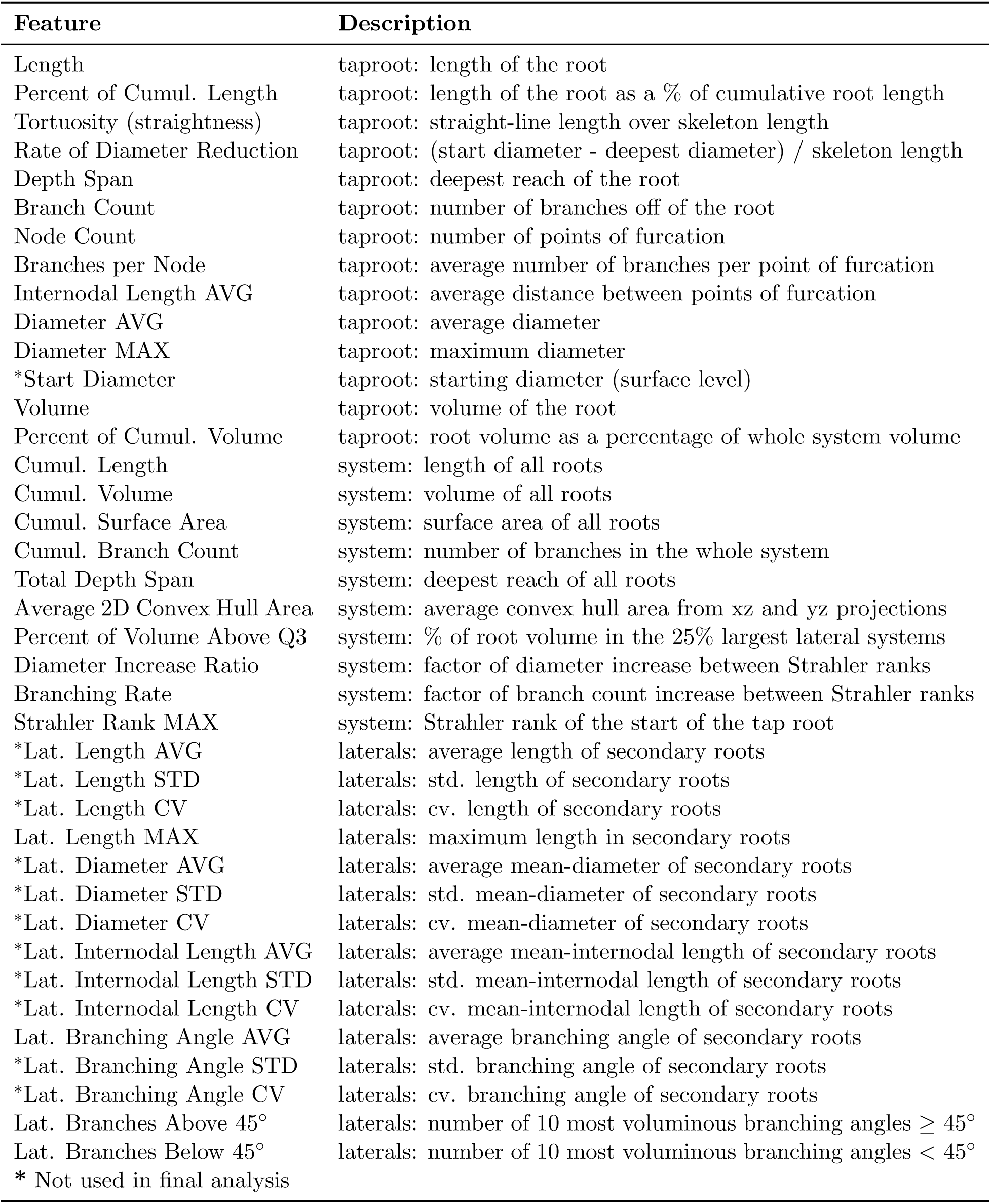
Features extracted from the point clouds of individual trees.

#### Soil Impact

Comparing RSA features across soil types reveals clear differences in root system development between clay and sandy loam substrates. Roots grown in sandy loam exhibit longer lengths, larger diameters, and greater variation in volume and branching patterns, suggesting a capacity for more efficient exploration and exploitation of heterogeneous soil environments. The low compaction in sandy loam likely allows root systems to expand adaptively, forming complex architectures optimized for local variation in water and nutrient availability. This flexibility may reflect divergent strategies: some plants develop minimal structures to meet resource needs, while others produce more extensive systems, both made possible by the looser soil conditions. In contrast, RSA in clay loam appears more constrained. Our results indicate reduced variability in structure, potentially limiting the spatial extent of root colonization. While early-season penetration depths are comparable between soil types, roots in clay loam are thinner initially and only thicken later in development. This delayed thickening may influence structural features such as tortuosity. Although prior studies [Popova et al., 2016] associate higher tortuosity with looser soils due to greater anchorage requirements, our findings suggest otherwise: clay-grown taproots exhibited greater tortuosity. We hypothesize that this results from mechanical constraints imposed by compacted soils. The slow thickening of the tap root that occurs in more compacted soil forces the roots to conform to local density variation, leading to more winding growth paths.

### 3.6 Limitations and Future Directions

While this study demonstrates the value of 3D reconstruction for quantifying root system architecture (RSA) and reveals clear soil-dependent RSA differences, several limitations should be acknowledged.

First, although the 3D models are geometrically accurate, there is a resolution limit imposed by the block size. This limitation disproportionately affects small-diameter lateral roots directly attached to the tap root. Block size decreases linearly, but close to the tap root, the linear tapering of the block size is still relatively large. So the fine-scale twists and turns of small-diameter roots close to the tap root may be smoothed out, potentially inflating their apparent diameter. A promising future improvement would be to use a non-linear block sizing scheme that adapts to local root diameter, allowing more precise modeling of fine structures.

A second technical limitation is the inability to disambiguate intersecting roots. Modeling roots in 3D reduces the occlusions and apparent crossings that are ubiquitous in 2D imagebased methods. However, when two roots physically touch, the 3D point cloud does not leave gaps between them, making it challenging to distinguish between the two roots. The skeletonization methodology employed here relies on detecting gaps between distinct structures, which is problematic when roots intersect. In this study, care was taken to pose the roots before data collection to reduce the number of intersections, but this artisanal approach reduces the throughput of our methodology. Future work will focus on developing automated strategies for intersection disambiguation to improve the completeness and accuracy of RSA skeletons.

Finally, although this study was designed to demonstrate the accuracy of the modeling workflow and the potential of high-fidelity 3D modeling for root system architecture (RSA) analysis, it is important to acknowledge the limitations of our experimental design. The observed differences in phenotypic response to soil texture were derived from only 30 RSA samples, evenly distributed across three timepoints and two soil types. The limited sample size reduces statistical power, particularly for detecting interaction effects between features. Although the broad trend of greater RSA plasticity in sandy loam compared to clay loam is supported by our companion study [Bogati et al., 2025], some of the finer patterns highlighted here may reflect sampling noise rather than true biological responses. This concern is especially relevant for LDA analysis, which is prone to overfitting when sample counts are low. Future studies with larger sample sizes will be important for validating these trends and strengthening our conclusions.

## 4 Conclusion

This study advances the analysis of root system architecture from 3D data by achieving three primary objectives. First, we introduced an open-source Python framework that enables automated quantification of field-grown RSA from point cloud models. Unlike prior approaches, our method is agnostic to point cloud acquisition modality and does not require assumptions about rooting direction or angle, producing highly accurate 3D models of naturally grown annual dicots. Second, we designed 39 extractable features to quantify the modeled RSA. Using hierarchical reconstructions of 30 soybean plants, we demonstrated our framework’s usability and identified 14 key RSA features sensitive to soil type variation.

Finally, we used these features to characterize soybean RSA responses to two distinct soil textures — sandy loam and clay loam. Root systems grown in sandy loam exhibited greater plasticity, with more variability in depth and branching angles, whereas those in clay loam displayed more constrained and uniform architectures. These findings illustrate the framework’s potential to uncover meaningful phenotypic variation in complex soil environments, providing a foundation for future field-based RSA analysis.

## A Appendix

**Figure A1:**
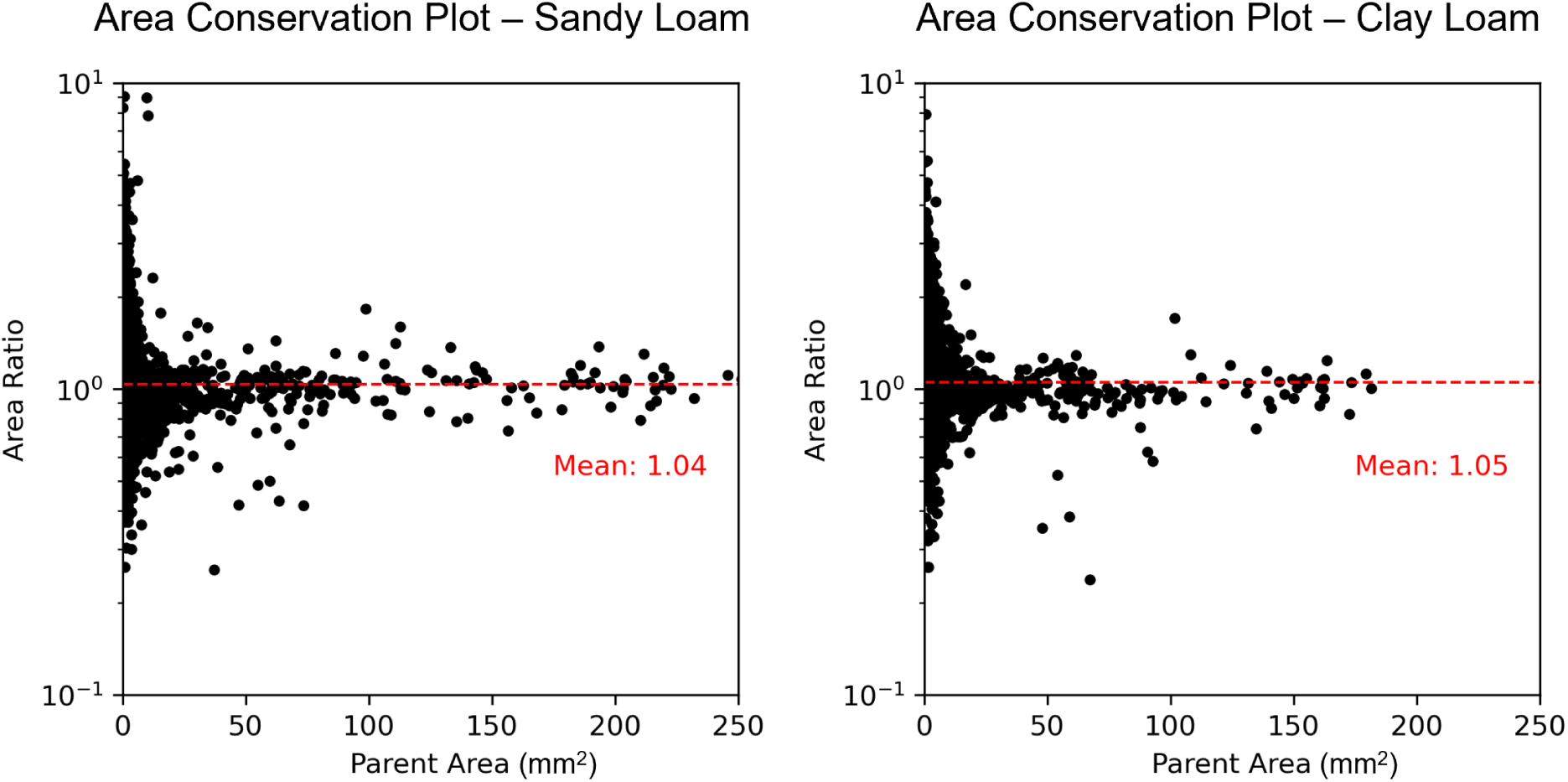
Area preservation analysis of roots grown in clay loam and roots grown in sandy loam. Mean area ratios (indicated by dotted red line) were calculated from segments have a diameter greater than 2mm. No significant difference existed across the two environments.

**Figure A2:**
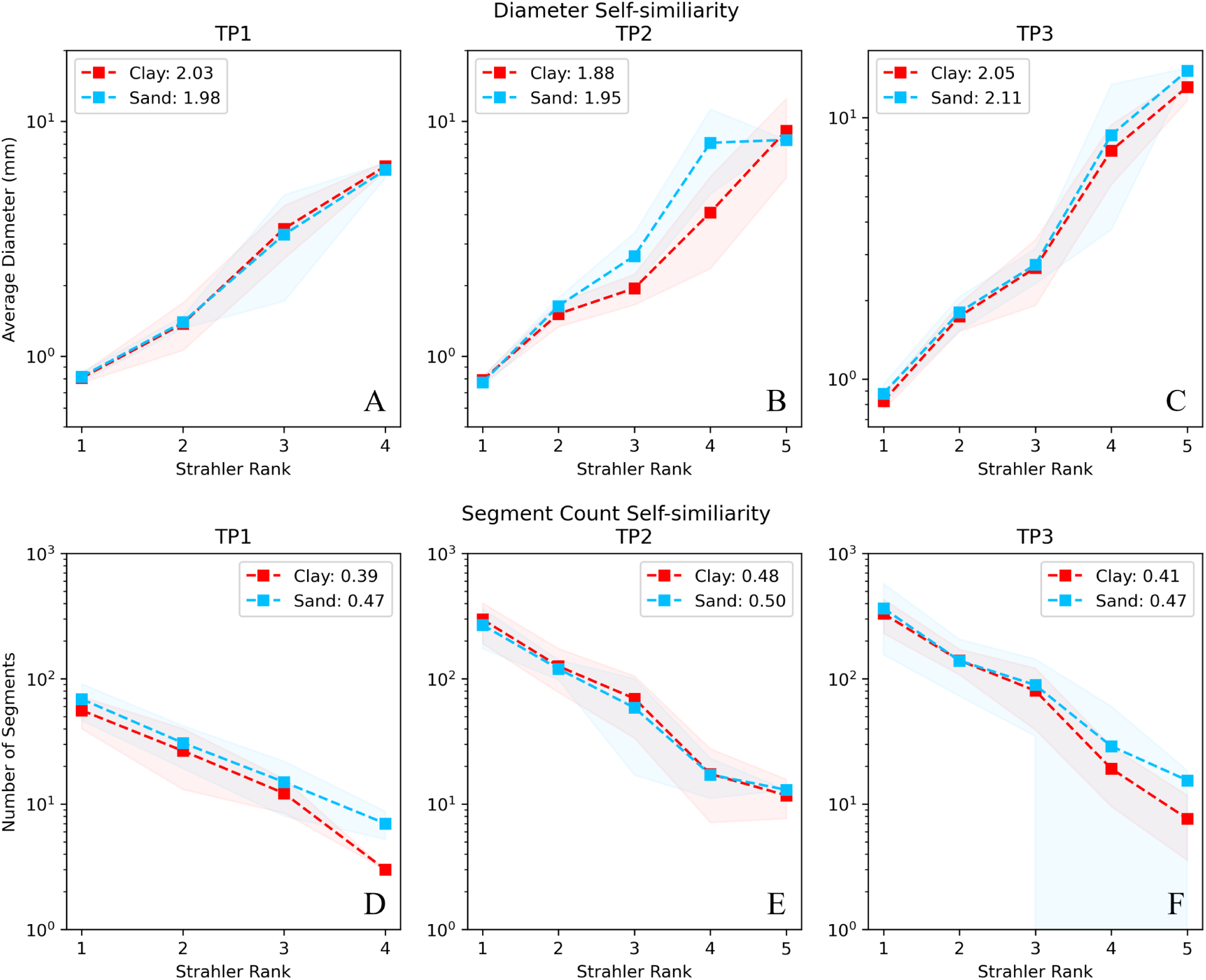
Self-similarity analysis of the diameter (A,B,C) and diameter (D,E,F) for each timepoint. The average diameter ratio *R_d_* and the branching ratio *R_n_* are shown in the plot legend. Shaded regions represent one standard deviation from the mean value. No significant difference was found in the branching rate within the two environments, suggesting that these two soils do not have a major effect on the self-similarity of the RSA.

**Figure A3:**
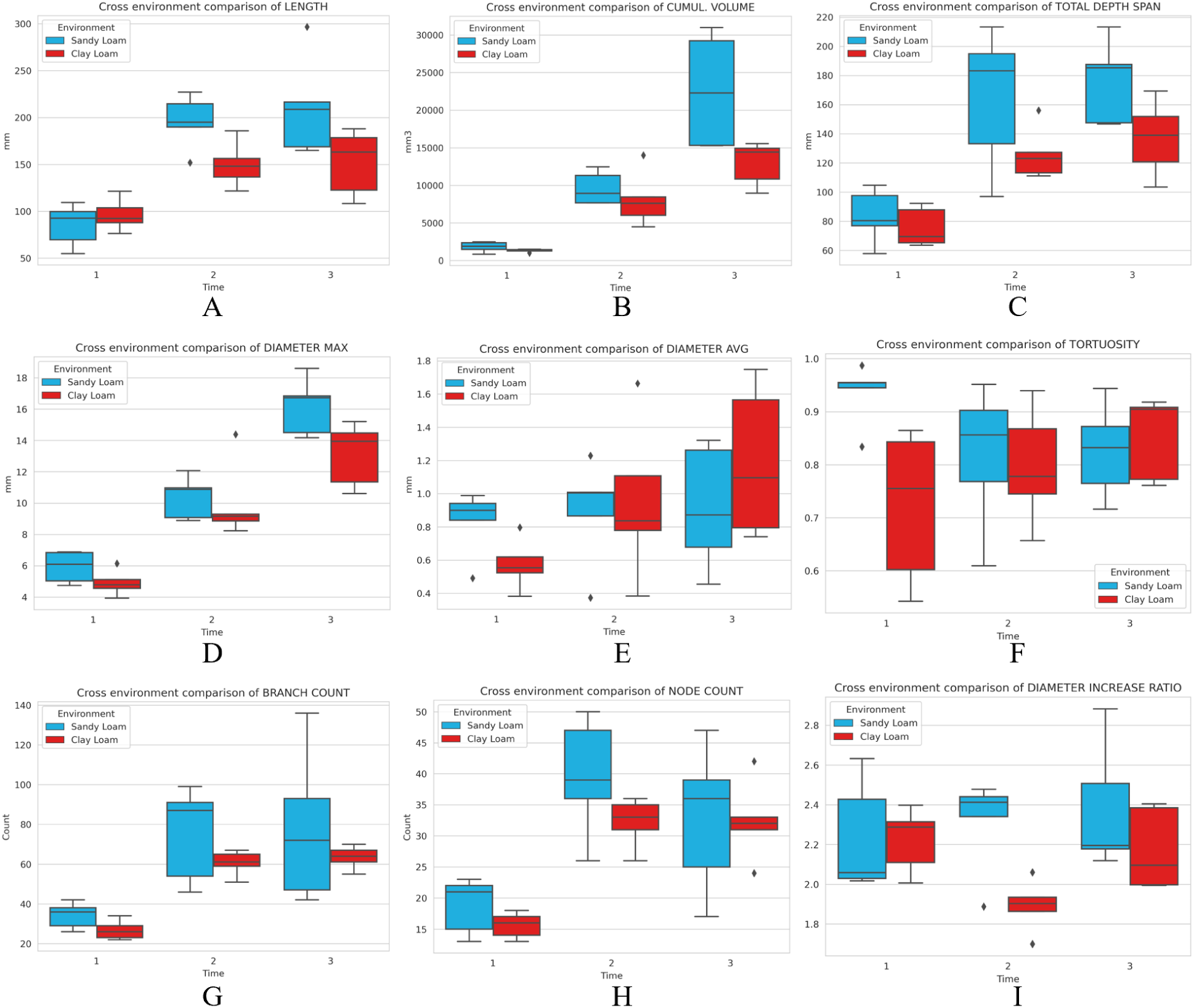
Box plots showing the distribution of select features by timepoint and environment. These indicate that sandy loam systems had longer (A), more voluminous (B), deeper reaching (C), and initially thicker (D) tap roots; while systems grown in clay loam showed shorter, stouter taproots, especially at late growing stages (E). Sandy loam systems showed far more variation in branch count (G), node count (H), and structural complexity (I); while clay loam systems had more tortuosity in root shape (F).

**Figure A4:**
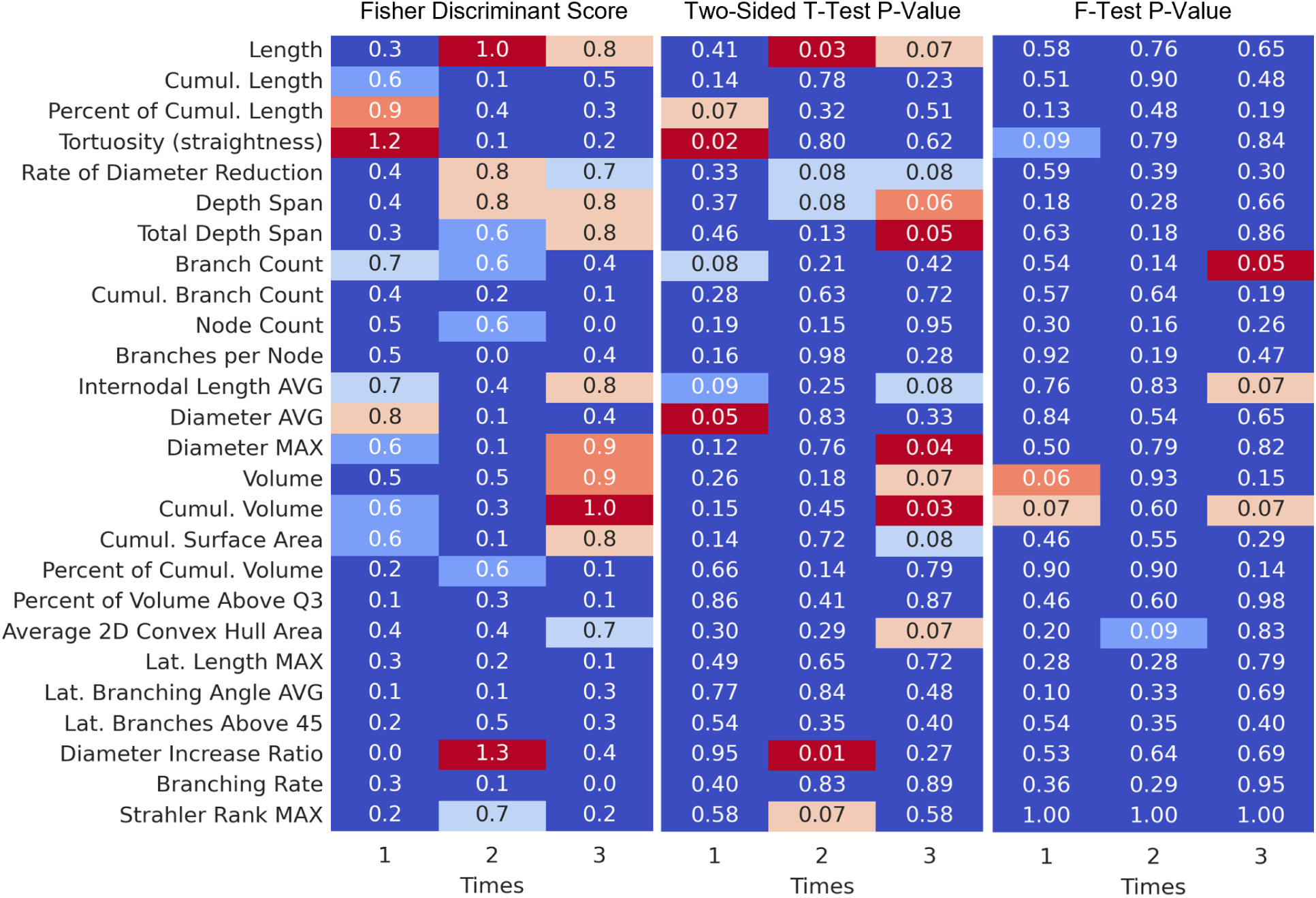
Feature distribution separability between clay and sandy loam is measured with fisher discriminant score. The statistical significance of the difference between the feature means from the two environments is quantified with the two-sided t-test. The statistical significance of the difference between the feature variations is measured using Levene’s F-Test. Generally, the RSA shows the most structural difference in timepoint 3 (late growth stage). Note that most of the distributions of the parameters describing the lateral roots showed no separability, significance, or difference in variance and were removed for brevity.

**Figure A5:**
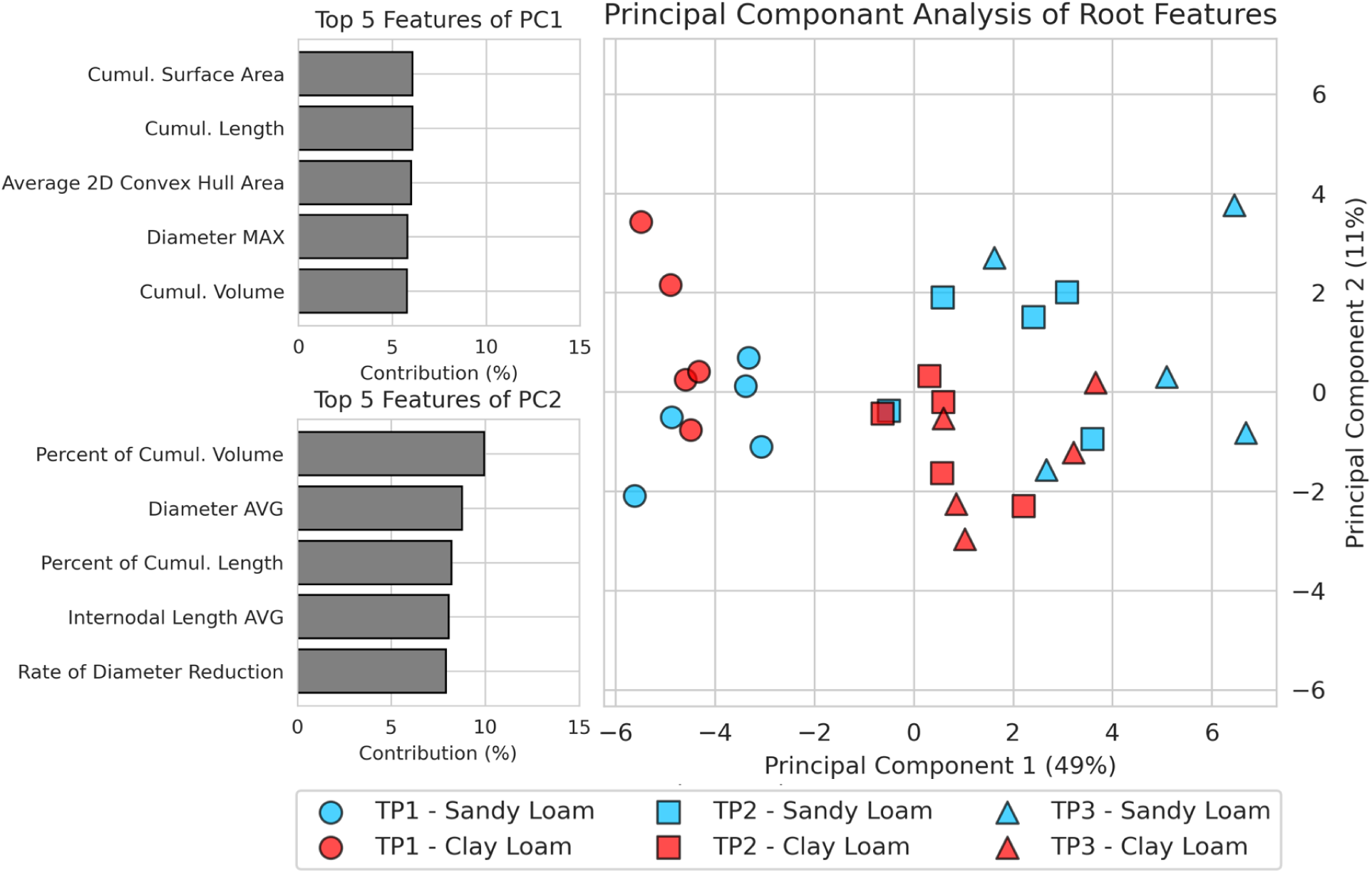
Comparison of features derived from 2D analysis and those derived from the 3D models. Red line is the best fit line. Panel A shows close agreement on the measures of Maximum Taproot Diameter. Panel B indicates that 2D measures tend to underestimate the lengths of lateral roots. Panel C shows a disparity between the average branching angle of lateral roots sampled from 2D imagery, and the average angle from the 3D RSA models.

**Figure A6:**
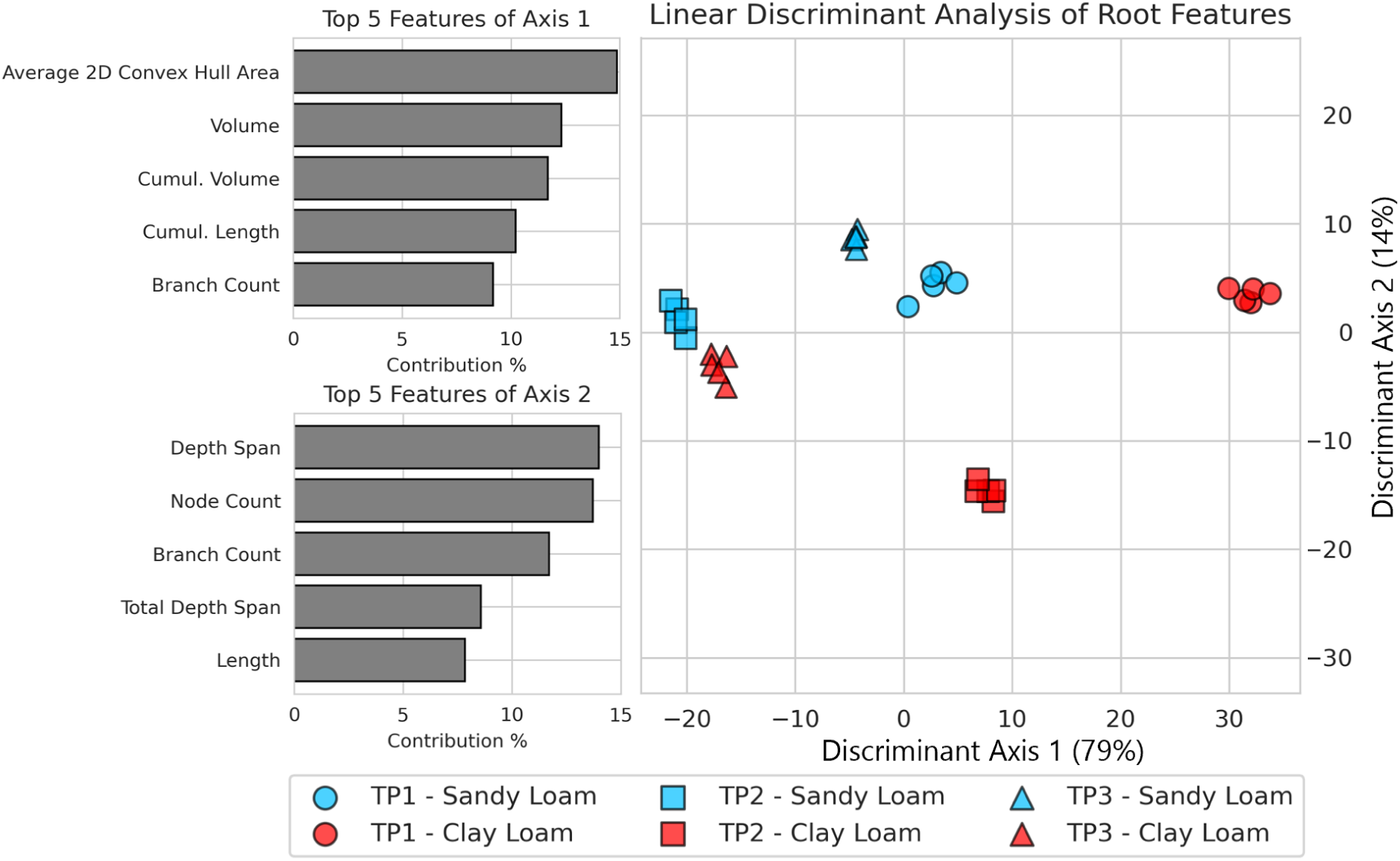
Comparison of the distributions of branching angles and lateral lengths. Feature samples from 2D images focus on major roots, leading to sample selection bias in some features. The top row (A) shows that the bias in 2D sampling tends to select more downward pointing roots, while the bottom row (B) indicates that short lateral roots are overlooked in 2D measurement. These two observations broadly demonstrate the difficulty in comparing 2D and 3D based metrics.

